# Cardioselective β-blocker Therapy Resolves *Staphylococcus aureus* Toxin-Induced Innate Immunoparalysis

**DOI:** 10.64898/2026.10.12.758599

**Authors:** Joshua W. Sun, Ruth Siew, Breanna Luna, David Gonzalez, Sachiko Kanaji, George Sakoulas

## Abstract

During *Staphylococcus aureus* sepsis, the pore-forming α-toxin (Hla) triggers lytic platelet death and provokes the release of catecholamines that modulate leukocyte effector function. The resulting surge in catecholaminergic action paralyzes host antimicrobial capacity by upregulating interleukin-10 (IL-10), profoundly suppressing the protective interferon (IFN) response to orchestrate innate immunoparalysis. Here, we deconvolute a multi-cellular regulatory pathway in the blood and highlight a pivotal immunological checkpoint, where pharmacological intervention with cardioselective β-blockers uncouples the immunosuppressive cascade, restores host antimicrobial function, and significantly improves survival outcomes.

## Main Text

*Staphylococcus aureus* is a ubiquitous commensal of the human microbiome, however under opportunistic circumstances, can emerge as the leading most causative agent of human sepsis (*1*). Despite improvements in modern supportive care, sepsis remains the most common cause of immediate hospital death (*2*). The invasive potential of *S. aureus* is largely attributed to the transcriptional accessory gene regulator, which controls virulent expression by regulating a number of cell wall-associated proteins, such as protein A, coagulase, fibronectin binding proteins, or by stimulating the release of the α, β, γ-exotoxins, leukotoxins, or TSST-1 (*3*). Among the most prominent is the pore-forming α**-** toxin (Hla). Secreted as a monomer, Hla oligomerizes into a β-barrel heptameric pore upon binding to its proteinaceous receptor ADAM10, allowing the flux of cations such as Ca^2+^ and K^+^ to initiate downstream aberrations and immune cell death (*4*). While sublytic concentrations of Hla elicit a silent cellular demise by apoptosis, higher membrane-perforating doses induce a contrastingly loud form of death by cellular lysis, known as pyroptosis or necroptosis (*5, 6, 7*).

Nonetheless, a standard function of inflammatory cell death is to prepare the host for microbial elimination and to promote the repair of damaged tissues (*8*). Excessive inflammation is, however, consequential, triggering a cytokine storm characterized by elevated levels of the pro-inflammatory cytokines IL-1, IL-6, TNF, IFN, IL-12, and IL-8 (*9*). This pathological inflammatory response can cause hypercoagulable events and microvascular thrombosis, while simultaneously consuming critical populations of circulating immune cells alongside their indispensable effector functions (*10*). Further is the induction of the archetypal anti-inflammatory IL-10 cytokine, which serves to counter-regulate the systemic inflammatory response, shifting the host toward an immunocompromised state (*11*). Maintaining the appropriate control of innate immune cell homeostasis is key to mitigating *S. aureus* bloodstream persistence and tissue dissemination. However, the mechanisms that drive these cellular processes remain ill defined.

Clearance of the pathogen while preventing collateral inflammatory damage presents a compelling challenge for the innate immune system. In our continued quest to identify host-directed therapeutic strategies to combat the rise of multi-drug resistant bacterial pathogens, we dissect the interaction between *S. aureus* and circulating innate immune cells to examine the underlying mechanisms of Hla-induced immunosuppression. Previously, we uncovered that *S. aureus* is highly susceptible to platelets, but resistant to leukocyte antimicrobial killing, and that Hla selectively causes thrombocytopenia to evade blood antimicrobial defense (*12*). Platelets vastly outnumber leukocytes in the bloodstream and constitute a primary reservoir for catecholamines. As such, we hypothesized that their lytic demise prompts a massive spillover of catecholamines into circulation, attenuating antimicrobial activity via autocrine and paracrine signaling.

We evaluated this hypothesis by examining the overall impact of catecholamines on human blood antimicrobial activity during live *S. aureus* infection. Whole blood isolated from healthy human donors treated *ex vivo* with the cardioselective β-blocker atenolol prior to *S. aureus* co-incubation significantly impaired bacterial survival (**Fig. 1a**). To parse the cellular drivers of this localized immunosuppressive phenotype, we evaluated the interactions of *S. aureus* Hla with isolated blood components. In platelet-rich plasma (PRP), infection with wild-type (WT) *S. aureus* demonstrated greater survival compared to the live attenuated isogenic Δ*Hla* mutant, an advantage that was significantly neutralized by metoprolol (**Fig. 1b**). Exposure of isolated human platelets to recombinant Hla provoked dose-dependent release of lactate dehydrogenase (LDH) and a concurrent release of norepinephrine (NE) (**Fig. 1c**), which was counteracted by β-blocker treatment (**Fig. 1d**). Because functional consequence upon β-adrenergic receptor activation elicits downstream elevation of Ca^2+^ and cyclic adenosine monophosphate (cAMP) to regulate cellular survival, we decided to measure these second messengers in isolated human platelets. Metoprolol was observed to subvert Hla-induced Ca^2+^ overload (**Fig. S1**), and dose-dependently protect platelets from Hla-induced lysis (**Fig. 1d**). Corresponding to cAMP as a negative regulator of innate immune cell effector function (*13*), directly elevating intracellular platelet cAMP levels with forskolin dose-dependently promoted bacterial survival (**Fig. 1e, 1f**), whereas metoprolol restricted bacterial survival (**Fig. 1g**).

**Fig. 1.**
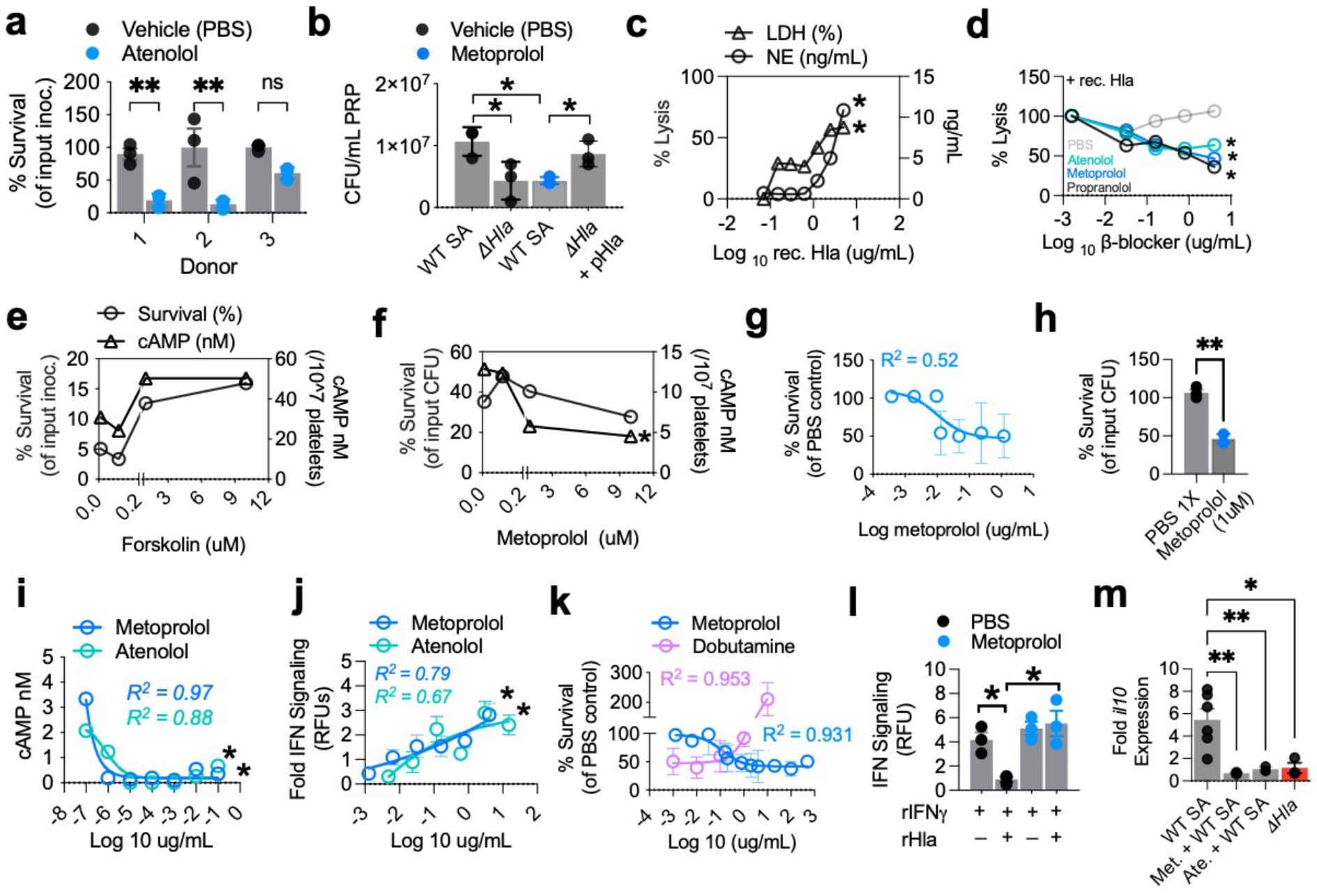
Cardioselective β-blockers enhance blood antimicrobial defense by blocking Hla-induced platelet death and IL-10 suppression of IFN signaling. **a**, Survival of WT *S. aureus* in human whole blood isolated from healthy donors following *ex vivo* pre-treatment with vehicle (PBS 1x) or 10 μm atenolol (MOI = 0.01). **b**, Survival of WT *S. aureus* compared to isogenic *ΔHla* mutant in human PRP in the absence or presence of 10 μm metoprolol. **c**, Cell death assessed by lactate dehydrogenase (LDH) release and concurrent extracellular norepinephrine (NE) release from isolated washed human platelets exposed to increasing concentrations of recombinant Hla. **d**, Isolated human platelets dose-dependently treated with metoprolol, atenolol, and propranolol followed by exposure to 5 μg/mL recombinant Hla. **e, f**, Quantification of intracellular cAMP levels in isolated human platelets treated with forskolin or metoprolol and co-incubated with WT S. aureus. **g**, Bacterial survival in isolated platelets dose-response treated metoprolol relative to PBS 1x control. **h**, Intracellular bacterial survival in purified human neutrophils pre-treated with PBS 1x or metoprolol *ex vivo* (MOI = 10). **i**, Intracellular cAMP concentrations in human neutrophils following WT *S. aureus* challenge with or without metoprolol and atenolol treatment. **j**, Relative leukocyte interferon (IFN) reporter signaling activity in human neutrophils challenged with WT *S. aureus* metoprolol. **k**, Bacterial survival in human monocytic THP-1 cells treated with escalating doses of metoprolol or dobutamine prior to WT *S. aureus* co-incubation. **l**, THP-1 Monocyte IFN reporter signaling following exposure to recombinant Hla with or without metoprolol rescue. **m**, RT-qPCR analysis of *il10* mRNA expression in THP-1 monocytes challenged with WT *S. aureus* or the Δ*Hla* mutant in the presence or absence of metoprolol or atenolol. Data are presented as mean ± s.d. or linear regression and is representative of at least 3 independent experiments. Statistical significance was calculated using a two-tailed unpaired Student’s t-test, one-way ANOVA with Sidak’s multiple comparisons test, or a non-linear regression; where applicable: *P<0.05, **P<0.01; ns, not significant. SA, *Staphylococcus aureus, MOI: Multiplicity of Infection, PBS: Phosphate buffered saline*

*S. aureus* can utilize host cells as intracellular reservoirs to protect against immune and antibiotic clearance (*14*). In our evaluation of Hla-induced intracellular Ca^2+^ signaling (**Fig. S1**), we observed that Ca^2+^ overload extends beyond that of platelets and therefore reasoned that the immunosuppressive β-adrenergic signaling pathway modulates neighboring immune cells as well. Freshly isolated human neutrophils pre-treated with metoprolol *ex vivo* prior to infection with WT *S. aureus* significantly restricted bacterial survival (**Fig. 1h**), correlating to a reduction in intracellular cAMP (**Fig. 1I**), and concurrently restoring protective IFN signaling (**Fig. 1J**). Similarly, metoprolol-treatment of human monocytic cells restricted WT *S. aureus* survival in contrast to dobutamine (**Fig. 1k**) and rescued IFN signaling (**Fig. 1l**). RT-qPCR analysis revealed that WT *S. aureus* significantly upregulated monocyte *IL10* mRNA expression compared to the isogenic Δ*Hla* mutant, a transcriptional induction that is markedly suppressed by metoprolol and atenolol (**Fig. 1m**).

We next asked if the localized immunosuppressive phenotype observed in tissue cell culture studies translates to infection outcomes *in vivo*. Outbred CD1 mice challenged intraperitoneally with a 50% lethal dose of WT *S. aureus* resulted in severe and rapid death compared to the live attenuated Δ*Hla* mutant (**Fig. 2a**). Systemic administration of the cardioselective β-blocker atenolol significantly enhanced host survival during WT infection. Utilizing a platelet depletion model, we observed that Hla requires platelets to induce cytokine storm, and that the targeted clearance of platelets from circulation completely subverts this toxicity, reducing IL-1β and IFN-γ cytokine levels to baseline states relative to Δ*Hla* (**Fig. 2b, c**). In platelet-depleted blood, we observed that Hla significantly inhibits IL-1β and IFN-γ levels, and interestingly, higher IFN-γ levels were detected in response to Δ*Hla* mutant when compared to the platelet abundant group (**Fig. 2b, c**). Together, this data reveals the dual nature of platelet immune function, where the response to WT *S. aureus* elicits both platelet-dependent cytokine storm yet mutes the immunogenicity of the live attenuated Δ*Hla*.

**Fig. 2.**
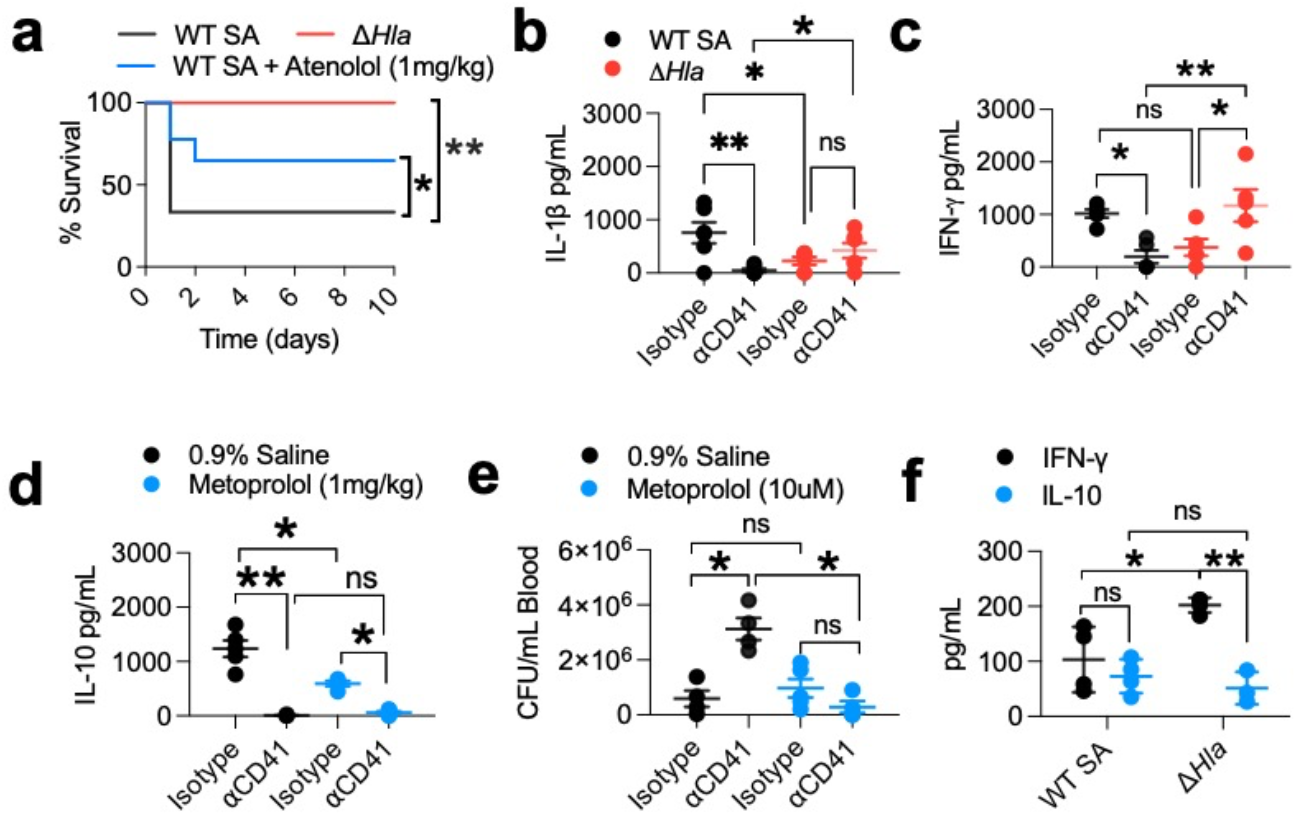
Cardioselective β-blockers improve survival outcomes and alleviates Hla-induced immunosuppression during murine sepsis. **a**, Kaplan–Meier survival curves of outbred CD1 mice challenged i.p. with WT *S. aureus* or the live-attenuated *ΔHla* mutant, with or without systemic administration of atenolol. **b, c, d** Concentrations of IL-1β, IFN-γ, and IL-10 in peritoneal exudates quantified 4 hours post-challenge in isotype IgG or ⍺CD41 treated mice infected with WT or the live attenuated *ΔHla* mutant. **d**, IL-10 concentrations measured in peritoneal exudate from isotype IgG treated or anti-CD41 treated mice challenged i.p. with WT *S. aureus* following treatment with 0.9% saline or metoprolol (n=5 /group, n=3 <LLD). **e**, *Ex vivo* bacterial burden in terminal cardiac blood harvested from isotype IgG control or ⍺-CD41 treated mice prior to 0.9% saline or metoprolol. **f**, Serum IL-10 and IFN-γ concentrations (n=5 /group, n=3 <LLD). Survival curves were analyzed using the log-rank Mantel–Cox test (*n =* 10 per group). Cytokine and CFU data are shown as mean ± s.e.m., and unless otherwise stated n= 5 mice/group). Statistical comparisons were performed using two-way ANOVA with Šídák’s multiple comparisons test; *\*P<0*.*05, **P<0*.*01; ns, not significant. SA, Staphylococcus aureus*

To investigate the hypothesis that sepsis lethality is driven by a concurrent surge in immunosuppressive β-adrenergic signaling, infection with WT *S. aureus* provoked a substantial release of IL-10, an event found to be entirely dependent upon the host platelet compartment, with a significant reduction observed in the metoprolol treated group (**Fig. 2d**). Blood harvested from platelet-depleted mice prior to treatment with metoprolol *ex vivo* significantly reduces WT *S. aureus* survival relative to the platelet abundant group (**Fig. 2e**). How systemic Hla toxicity actively cripples protective blood immunity during intravenous infection was determined and IFN-γ was detected at higher levels in response to *ΔHla* mutant relative to the WT parent (**Fig. 2f**).

In sum, we describe a mechanism by which Hla hijacks β-adrenergic signaling to deprive the host of mounting a proper immune response. A growing body of evidence demonstrates that cardioselective β-blockade improves sepsis mortality outcomes (*15, 16*), and despite the seemingly counterintuitive use of a negative inotropic agent, we often ask at what cost to host immunity are we supporting cardiac function? Suppose the exogenous administration of norepinephrine compounds the endogenous sympathetic overdrive and cascades the host toward a cyclical phase of immunoparalysis and recurrent infections (*17, 18*). Perhaps it’s entirely conceivable, then, that intervention during post-sepsis recovery may upend this vicious cycle. Consistent with prior studies (*19*), this paradigm reflects the underlying biological architecture of *S. aureus* immune evasion and highlights a fundamental immunological checkpoint for the safe and efficacious design of host-directed therapeutics. It follows, therefore, that any prophylactic measure made against *S. aureus* Hla should aim to circumvent the catecholamine dump, or risk severing the connective layer between innate recognition and adaptive memory.

## Materials and Methods

### Ethics Statement

Human whole blood was obtained via venipuncture from healthy volunteers under written informed consent approved by the UC San Diego Human Research Protection Program and Institutional Review Board. Animal studies were conducted in accordance with protocols approved by the UC San Diego IACUC. All efforts were made to minimize animal numbers and suffering.

### Bacterial Strains and Growth Conditions

The live attenuated isogenic *Δ*Hla knockout strain was constructed from the USA300 strain by precise allelic replacement of the encoding *hla* gene. WT USA300 and the isogenic *ΔHla* mutant strains were grown overnight, shaking at 37°C in Todd-Hewitt broth. Following overnight growth, bacterial cultures were centrifuged, washed once in 1x phosphate-buffered saline, and resuspended to the required optical density to achieve the targeted multiplicity of infection for *in vitro* assays or colony forming unit (CFU) concentration for *in vivo* challenge.

### Human Platelet and Neutrophil Isolation

Human whole blood was drawn with a 25-gauge needle into tubes containing Anticoagulant Citrate Dextrose Solution A (ACD). To obtain PRP, whole blood was centrifuged at 100 × g for 10 min without braking. For washed platelet assays, PRP was further centrifuged in the presence of prostacyclin to prevent premature activation, washed, and resuspended in ACD buffer.

For neutrophil isolation, whole blood collected in sodium heparin tubes was carefully layered over a standard polymorphonuclear cell density gradient medium and centrifuged according to the manufacturer’s instructions. The polymorphonuclear band was extracted, washed with HBSS without Ca^2+^/Mg^2+^, subjected to hypotonic erythrocyte lysis, and resuspended in RPMI 1640 medium supplemented with 2% heat-inactivated fetal bovine serum.

### THP-1 Monocyte Tissue Culture

The human monocytic leukemia cell line THP-1 was cultured and maintained in RPMI 1640 medium supplemented with 10% heat-inactivated FBS, 1% penicillin-streptomycin, and 2 mM L-glutamine. Cells were maintained in a humidified incubator at 37°C with a 5% CO_2_ atmosphere. Prior to infection assays, THP-1 cells were centrifuged, washed to remove antibiotics, and seeded into tissue culture plates in antibiotic- and serum free assay medium.

### *In Vitro* and *Ex Vivo* Infection Assays

For all *in vitro* and *ex vivo* infection studies, host cells were incubated with *S. aureus* at 37°C for 2 hours in a 5% CO_2_ atmosphere.

#### Platelet and Whole Blood Assays

Human whole blood, PRP, or isolated washed platelets were pre-treated for 15 min with phosphate buffered saline or the cardioselective beta-blockers atenolol or metoprolol. Samples were then inoculated with *S. aureus* at an MOI of 0.01 for 90 minutes. To enumerate surviving bacteria, samples were sonicated to disrupt host cells, serially diluted in molecular grade H_2_O, and plated on Todd-Hewitt agar. Platelet lysis was quantified via lactate dehydrogenase release assays following exposure to recombinant Hla. Intracellular Ca^2+^ mobilization was measured every 30 seconds for up to 30 minutes by loading platelets with 2 µM Fluo-3 dye and tracking fluorescence upon Hla stimulation.

#### Leukocyte Assays

Isolated human neutrophils and THP-1 monocytes were pre-treated with vehicle, atenolol, or metoprolol for 2 hours and subsequently infected with *S. aureus* at an MOI of 10 for 2 hours at 37°C. Bacterial survival was enumerated by serial dilution and plating on THA. Intracellular cAMP levels in both platelets and leukocytes were quantified using a competitive cAMP ELISA. Leukocyte interferon signaling capacity was measured using targeted fluorescent reporter assays. To evaluate immunosuppressive IL-10 cytokine production, THP-1 cells were lysed post-infection, RNA was extracted, and *il10* expression was quantified by RT-qPCR relative to the *ΔHla* mutant baseline.

### *In Vivo* Sepsis Model

Outbred 8- to 10-week-old CD1 mice were utilized for *in vivo* sepsis modeling. Mice were challenged via intraperitoneal injection with a 50% lethal dose of WT *S. aureus* or the live-attenuated isogenic *ΔHla* mutant. Survival outcomes were recorded over a 10-day observation period. Mice that appeared moribund were euthanized by CO_2_ asphyxiation.

For platelet-depletion models, mice were treated with an ⍺CD41 antibody for 16 hours prior to infection, reducing circulating platelets to approximately 15% of baseline abundance. To assess systemic inflammation and immune paralysis, peritoneal lavage fluid and blood was collected by terminal cardiac puncture 24 hours post-challenge. Concentrations of IL-1β, IL-10, and IFNγ were quantified using ELISA.

## Acknowledgements

We thank George Y. Liu from the Department of Pediatrics at UCSD and Rady’s Children’s Hospital for insightful discussions. This work was supported by the T32 Molecular and Cellular Biology of Allergy Training Grant (5T32AI007469-29).

## Author contributions

J.W.S. conceived the study, performed experiments, analyzed the results, and wrote the manuscript with conceptual and structural input from all co-authors.

## Supplementary materials

**Supplementary Fig. 1.**
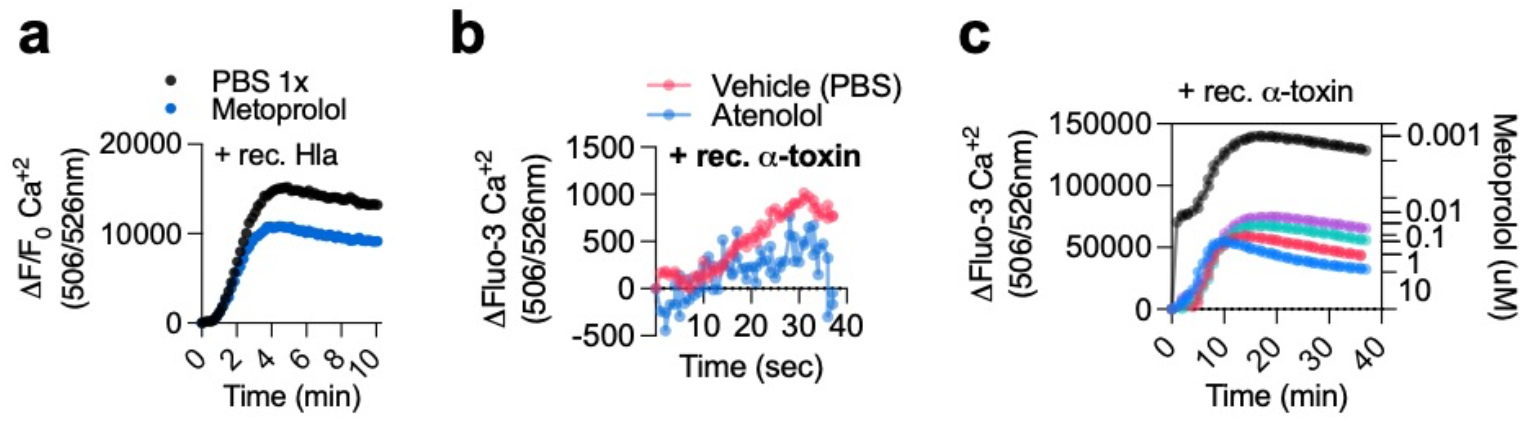
Cardioselective β-blocker treatment attenuates Hla-induced Ca^2+^ mobilization. Kinetic traces of intracellular Ca^2+^ mobilization in fluo-3-loaded dye in isolated human **a**, platelets **b**, neutrophils, or **c**, THP-1 monocytes stimulated with 1µg/mL recombinant Hla in the presence of vehicle or metoprolol, recorded every 30 seconds by fluorescence. Data representative of at least n>3 independent experiments.

